# The overlooked first intercostal ligament: Does it help to stabilize the Weberian apparatus?

**DOI:** 10.1101/2023.11.20.567829

**Authors:** Jake Leyhr, Tatjana Haitina, Nathan C. Bird

## Abstract

The Weberian apparatus is a novel hearing adaptation that facilitates increased hearing sensitivity in otophysan fishes. The apparatus is a complex system composed of modifications to anterior vertebral elements, the inner ear, and the swim bladder. A critical piece of the system that often receives minor attention are the various ligaments that bridge these three regions. Most famous of the ligaments is the interossicular ligament, which connects the weberian ossicle chain (scaphium-intercalarium-tripus). Several other ligaments are present, including the suspensor (tripus to parapophysis 4) and the triple ligament (tripus-os suspensorium-tunica externa). Here, by combining diffusible iodine-based contrast enhancement (DICE) and propagation phase-contrast synchrotron radiation micro-computed tomography (PPC-SRμCT) with classic histological methods we shine new light on the first intercostal ligament (ICL1) and discuss its potential function in relation to the Weberian apparatus. ICL1 is nearly absent from the cypriniform literature, typically only mentioned in a general discussion together with other intercostal ligaments. This study examines the development and structure of ICL1 comparatively with the other definitive Weberian ligaments in the zebrafish (*Danio rerio*). We provide a comprehensive view of development, three-dimensional shape, and composition to generate hypotheses regarding potential functions of ICL1 within the greater Weberian apparatus. Given new detail presented herein regarding the structure of ICL1, modifications to rib 5 and parapophysis 4 for ICL1 attachment, and the alignment of ICL1 with the os suspensorium, we propose a supportive (anchoring) role of ICL1 to aid in minimizing non-optimal movement of the structures of the fourth vertebra, thereby allowing the focus of vibrations anteriorly to the ossicle chain with minimal signal loss in zebrafish and other species with similar Weberian apparatus morphologies. We conclude that ICL1 should be included in future analyses of Weberian apparatus function where ligaments are addressed.

## 1. INTRODUCTION

The Weberian apparatus is a complex and novel morphological system used to enhance hearing in a diverse collection of fishes and unites the more than 10,000 species in Series Otophysi, the largest group of primarily freshwater fishes (Nelson et al. 2016). The bony and soft tissue components of the Weberian apparatus vary substantially among species, both phylogenetically among the orders (Cypriniformes, Characiformes, Siluriformes, and Gymnotiformes), as well as within orders in various environmental systems, such as within Cypriniformes (Alexander 1962; Bird & Hernandez, 2007; Bird et al. 2020a) and Siluriformes (reviewed in Ladich, 2023). The morphological variation is likely intimately tied to unique noise-regimes created in these different environments, and various modifications in both hard and soft tissues (Bird et al., 2020a) are employed to maintain optimal functionality as evidenced by compromised function when shifting to different environments (Amoser & Ladich, 2005), however some plasticity has been found among cypriniforms in different environments (Ladich, 1999).

The Weberian apparatus consists of several modifications, including to the inner ear, swim bladder, and anterior vertebrae. The most famous elements are the Weberian ossicles (claustrum, scaphium, intercalarium, tripus, and os suspensorium), and are easily identified and examined in the relatively simple Weberian apparatus morphology seen in most cyprinids, such as the zebrafish (Bird & Mabee, 2003). The scaphium, intercalarium, and tripus act as a chain for direct transmission of vibrations generated by the swim bladder (in response to sound) to the sinus impar (then into the ear) via ligamentous attachments (Figure 1). The vertebral elements of the Weberian apparatus are modifications of several skeletal elements, including neural arches (scaphium, intercalarium) and parapophyses and ribs (tripus, os suspensorium). While the claustrum and os suspensorium are not generally attributed a primary role in Weberian apparatus function, their intimate association with the ossicle chain and the sinus impar (claustrum) and swim bladder (os suspensorium) suggests a supportive or stabilization role, rather than transmissive. Collectively, the apparatus transforms pressure waves collected by the swim bladder and transmits and likely amplifies them to the inner ear where they can be detected by differential motion in the otolith-based inner ear (Ladich & Popper, 2004; Ladich & Schulz-Mirbach, 2016). This allows for detection of wider frequency ranges as well as sensitivity at lower sound pressures compared to species with non-modified hearing (Fay, 1988; Fay and Simmons, 1999; Schellert and Popper, 1992).

**Figure 1:**
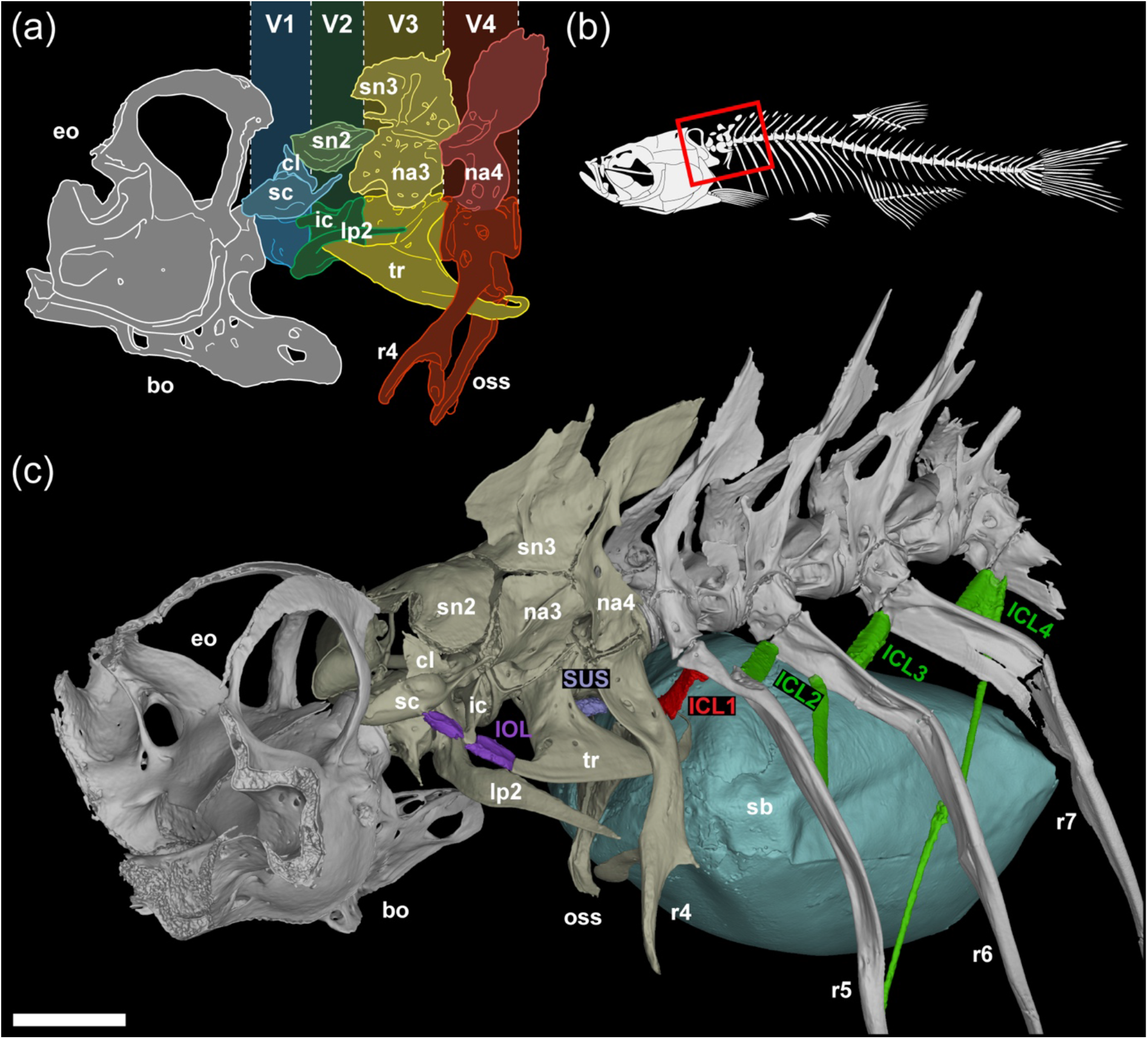
(a) Schematic of the adult zebrafish Weberian apparatus (cervical vertebrae 1-4) and occiput. (b) Schematic of zebrafish skeleton with red box highlighting the region containing the occiput, Weberian apparatus, and anterior rib-bearing vertebrae displayed in (c). (c) 3D rending of 24mm SL zebrafish Weberian apparatus in association with the occiput, swim bladder, and rib-bearing vertebrae, with ossicular and intercostal ligaments highlighted in purple, lilac, red, and green. Ventrally-projecting bones from the right-hand side of the vertebrae were removed to simplify the visualisation. bo: basioccipital, cl: claustrum, eo: exoccipital, ICL: intercostal ligament, IOL: interossicular ligament, lp: lateral process, na: neural arch, oss: os suspensorium, r: rib, sb: swim bladder, sc: scaphium, sn: supraneural, SUS: suspensor ligament, tr: tripus, V: vertebra. Scale bar = 500µm.

### Weberian Apparatus Studies in Cypriniform Fishes

#### Evolutionary variation

A rich lineage of descriptive research is found for the Weberian apparatus, especially for catfishes, dating back to Weber’s initial description of *Silurus glanis* (1819,1820; see Ladich, 2023 for historical review). The research history for cypriniform fishes is nearly as long, and work on evolutionary variation, development, function, and genetics are available for both groups. For simplicity, we will focus primarily on the work done with cypriniform fishes. Within the vertebral region of the Weberian apparatus, substantial morphological research has focused on the morphology and evolution of the ossicles and other skeletal elements of the Weberian apparatus, including various bony encapsulations of the swim bladder (Ladich, 2023; Bird et al. 2020a). On the other hand, comparatively limited research has examined the structure, development, and evolution of the various ligamentous components within the vertebral region.

#### Ontogenetic studies

Several developmental/ontogenetic studies on the vertebral portion of the cypriniform Weberian apparatus have been conducted on multiple cypriniform species. Detailed studies on ontogeny in the zebrafish can be found for each region of the Weberian apparatus, including the inner ear (Kimmel et al., 1995; Haddon & Lewis, 1996; Bang et al., 2001; Bever & Fekete, 2002; Wang et al. 2015), vertebrae and ossicles (Bird & Mabee, 2003; Grande & Young, 2004), and swim bladder (Finney et al., 2006; Robertson et al., 2007; Winata et al., 2009). Ontogenetic study of the complete Weberian apparatus is also available for zebrafish (Bird et al., 2020a).

#### Genetic studies

Owing to its role as a model species, several genetic studies on the zebrafish have been published. These studies have generally focused on one of the regions. Classic mutant analyses have been available for years on the inner ear (Malicki et al., 1996; Whitfield et al., 1996) and swim bladder (Winata et al., 2009).

While the genetics of general body development, somitogenesis, and somite differentiation have been studied in detail in zebrafish, often the studies end before enough skeletal development in the axial skeleton has occurred to determine whether loss of a particular gene affects the ossicle chain or other vertebral-based elements. The role of *tbx6*, a critical gene regulating segmentation, has been examined in zebrafish (Akama et al. 2020). In *tbx6* null mutants, severe pattern-based defects are found in the axial skeleton, particularly severe in non-Weberian vertebral levels (Akama et al., 2020). In contrast, effects in Weberian vertebrae appear less severe, which aligns with the limited defects seen in the patterning of presumptive Weberian somites with regard to muscle fiber and dermomyotome patterning when compared to more posterior somites (Windner et al. 2012). Recently, the role of *nkx3.2* in Weberian apparatus development was examined, with null mutants revealing substantial defects in the axial skeleton (Waldmann et al., 2021) as well as in both the mineralized skeletal and soft tissue elements of the vertebral region of the Weberian apparatus (Leyhr et al. 2023).

Limited hormonal regulation has been examined as well, centering on thyroid hormone, which found varied morphological responses depending on level of thyroid hormone perturbation in various cyprinids (Kapitanova & Shkil, 2014) including zebrafish (Keer et al., 2019).

#### Previous study focus

Within the vertebral region, most of the focus has tended to be on the skeletal components, with limited discussion regarding the ligaments that connect them. Indeed, the ligaments within this region play a pivotal role in proper transmission of motion between the ossicles, and therefore the function of the apparatus as a whole. Several ligaments can be found, the most often referenced being the interossicular ligament within the ossicle chain. Several others are also found, though, and these ligaments vary substantially in fiber density, cellularity, and fiber composition, suggesting a variety of functional roles within the system (Bird et al., 2020a).

### Review of Ligaments and Past Ligament Research

Within the Weberian apparatus, four confirmed ligaments have been discussed within the literature to various degrees of depth: Interossicular (anterior and posterior), Suspensor, and Triple ligaments. We propose a potential fifth ligament, Intercostal 1 (between Rib 4 and Rib 5), merits additional attention based on its location, attachment, composition, and the likely role of intercostal ligaments in the evolution of the Weberian apparatus. Each of these ligaments are reviewed below.

#### Interossicular (anterior and posterior)

The most obvious and critical ligament in the system is the interossicular ligament (IOL), which ties the anterior process of the tripus to the manubrium of the intercalarium (IOL posterior), and the manubrium to the lateral surface of the scaphium (IOL anterior). The interossicular ligament, therefore, plays a direct role in sound transmission in the apparatus, is a key component for functionality, and is typically described or mentioned in most species-level descriptions of the Weberian apparatus. The interossicular ligament has a particular histological signature, being fiber dense and likely high tensile strength. Within the literature, it is included most often in anatomical descriptions (sometimes the only ligament mentioned).

While nearly uniformly referred to as a ligament, Chranilov (1927) referred to the IOL as a tendon. The IOL has tense collagen fibers identified by multiple stains, including Hall Brundt’s Quadruple Stain and Mallory’s triple stain. Alexander (1962) also noted that the IOL contains ichthyocol (most often associated with the swim bladder), which is collagen arranged morphologically to appear like short needles. While likely a form of collagen I, the precise relationship of ichthyocol to the various collagens has not been analyzed at the molecular level.

#### Suspensor ligament

Another large ligament is the suspensor ligament, which attaches the base of the tripus to the base of parapophysis 4 (which bears rib 4 and the os suspensorium). This ligament tends to be much less fiber dense (Alexander, 1962) and much higher in cellularity (Bird et al. 2020a,b; named the tripus-parapophysis 4 ligament). Based on chemical staining, Alexander (1962) described this ligament as composed of elastin, with collagen absent, making it starkly different from the IOL; however, no molecular analyses are available that could support this. While its specific role is unclear, it likely plays a supportive role in limiting the rocking of these elements in specific planes, possibly to help minimize signal loss during sound transmission.

Several studies have mentioned the suspensor ligament with various levels of description. Nelson (1948) described the suspensor in Catostomidae (however he did not explicitly name it) and postulated that it kept the tripus in place along with the triple ligament. Chardon and Vandewalle (1997) mention a “suspensor” ligament, however based on location, connectivity, shape (thick and crescent-shaped) and composition (collagen), they likely described the triple ligament. The suspensor ligament has also been described in *Gobio* (Vandewalle, 1974), *Barbus* (Vandewalle, 1989), zebrafish (Bird et al, 2020b; Leyhr et al., 2023), *Botia* (Alexander, 1964a) and more broadly across cypriniforms (Alexander, 1962; Bird et al. 2020a).

#### Triple Ligament

The triple ligament is a small, usually fan-shaped ligament that connects the transformator process of the tripus, os suspensorium, and swim bladder (Bird et al. 2020a,b). Some early descriptions in cyprinids (Evans, 1925) and *Hybognathus* (Niazi & Moore, 1962) mistook this ligament for a muscle, hence the alternate name of ‘tensor tympani’ in early literature, and may have led to several instances of the ligament being described but unnamed in the mid-1900s. Nelson (1948) described it (unnamed) as having a role in maintaining the position of the tripus. Mookerjee et al. (1952) described it (also unnamed) as a band of connective tissue connecting the os suspensorium and the transformator process in *Esomus*. Rojo (1987) also described the ligament in *Barbus* but left it unnamed.

The triple ligament in *Danio* was described by Bird et al. (2020b) as a dense, collagen rich ligament, roughly fan-shaped. Bird et al. (2020a) also described it in *Danio*, *Gyrinocheilus*, and *Ambastia*, but left it unnamed. A similar description was found by Alexander (1962; Ligament 1), who described it as fan-shaped bundles of radiating collagen fibers. Chardon and Vandewalle (1997) postulated that the triple ligament was a specialized intercostal ligament under the assumption that the os suspensorium is a modified Rib 4 while the transformator process is a modified Rib 3.

#### Intercostal 1 (intercostals overall)

Intercostal ligaments are most often omitted from descriptions of the Weberian apparatus, due to the focus on the anterior-most vertebrae (1-4). When present, they are typically drawn in figures but not discussed in the text. An interesting variety of intercostal ligament patterns and lengths are found in many species, including cyprinids (Liao & Kullander, 2013). Many questions still remain regarding the origin and homologies of both the skeletal elements of the Weberian apparatus and its ligaments. Rosen and Greenwood (1970) hypothesized that the interossicular ligament was likely homologous to ancestral inter-neural arch ligaments (IOL anterior, scaphium to intercalarium) and a combination of inter-neural arch and neural arch-parapophysis ligaments (IOL posterior; intercalarium to tripus). This would make sense given the consensus homologies of the scaphium (neural arch 1), intercalarium (neural arch 2), and tripus (largely parapophysis 3), however questions remain regarding the ubiquity of these types of ligaments in most fishes. Unfortunately, they did not address the suspensor or triple ligaments, but based on their drawings one could make a strong claim that their intercostal ligament between R3 and R4 would be the precursor to the suspensor ligament. Intercostal ligaments were explicitly drawn by Chardon and Vandewalle (1997), including between R4 and R5. Following the initial suggestion of Sagemehl (1885), they hypothesize that the IOL represents the transformation of a swim bladder outgrowth, similar to those found in clupeids, into an elongated ligament. Their hypothetical ancestral reconstruction has a broad, dense collagenous ligament extending between the tripus and parapophysis/rib 4, suggesting an intercostal origin for at least the suspensor ligament, in agreement with Alexander (1962).

The first intercostal ligament is found bridging and uniting the head of rib 5 to the region where rib 4 and the os suspensorium split on vertebra 4, based on position and connectivity this ligament could provide specific support to the structures of vertebra 4, limiting their movement (likely restricting dorsal-ventral flexion or rotation at the centrum). The intimate association of parapophysis/rib 4 and parapophysis/rib 5 has been noted in other cypriniform species (Conway, 2011). For this study, we re-examine this ligament and compare the structure and development of the first intercostal ligament with the other main Weberian ligaments to form hypotheses regarding its possible functional role within the system. Similar histological signatures, specific insertion sites, and developmental timing may provide evidence of the importance of this ligament, as well as a broader role of vertebra 5 in the simplified zebrafish Weberian system (generally considered limited to V1 through V4). Taken with other morphological elements of vertebra 5, a reexamination of this vertebra in the sphere of Weberian apparatus evolution is warranted, as it may represent a transitional vertebra that develops, at least in part, within the Weberian apparatus developmental module and play a functional role within the Weberian system.

## 2. MATERIALS AND METHODS

### 2.1 DICE-PPC-SRµCT

Juvenile zebrafish (*Danio rerio*, AB line) were obtained at 1mpf and 2mpf (8.5mm and 24.0mm SL, respectively). Following euthanasia using an overdose of MS-222 (300mg/L) and fixation in 4% paraformaldehyde, diffusible iodine-based contrast enhancement and propagation phase-contrast synchrotron radiation micro-computed tomography (DICE-PPC-SRµCT) was performed as previously described by Leyhr et al. (2023), at BM05 of the European Synchrotron Radiation Facility - Extremely Brilliant Source (ESRF-EBS) in France, and with voxel sizes of 0.727µm and 3µm. Reconstructed jpeg2000 image stacks were imported into VGStudio MAX (v3.5.1) for manual segmentation and rendering.

All animal experimental procedures were approved by the local ethics committee for animal research in Uppsala, Sweden (permit number 5.8.18-18096/2019). All procedures for the experiments were performed in accordance with the animal welfare guidelines of the Swedish National Board for Laboratory Animals.

### 2.2 Histology

Juvenile and adult zebrafish (*Danio rerio*, AB line, 5.5-33.7mm TL) were obtained from the Zebrafish International Resource Center (Eugene, Oregon). Adults were anesthetized using buffered 0.04% MS-222, fixed in chilled 10% buffered formalin (24h at 4°C minimum). Three specimens were paraffin-embedded and sectioned at 8mm (one specimen per plane – sagittal, horizontal, and coronal) and processed as previously described (Bird et al. 2020a). Sections were stained using a modified Hall-Brunt Quadruple stain (Hall, 1986), allowing for visualization of ligaments as well as bone, cartilage, and ossification sites within cartilages.

Histological images were collected using a Zeiss Axio ScopeA.1 microscope with a ProgRes CF Scan camera and CapturePro v2.8.8 software (Jenoptik). When necessary, composite histological images were assembled in Adobe Photoshop 2022 using the Automate-Photomerge feature (default settings) to remove stitching errors.

All histological-based procedures followed an approved University of Northern Iowa IACUC protocol (#2015-05-1). All tissues are deposited in Biological Collections in the University of Northern Iowa Department of Biology.

## 3. RESULTS

### 3.1 Structural comparison of intercostal ligament 1 to Weberian ligaments and posterior intercostal ligaments

The first intercostal ligament (ICL1), connecting rib 5 to parapophysis (rib) 4 has similarities to both Weberian ligaments and more posterior intercostal ligaments, as well as unique features bridging the different sets of ligaments (Figure 1c). The Weberian ligaments vary the most in composition, with the suspensor ligament (SUS, Figure 2, middle row) having a markedly different histological signature (elastin) compared to the strong collagen 1 staining seen in the interossicular ligament (IOL, Figure 2, top row) and the triple ligament (not shown). ICL1 is remarkably similar to the IOL (Figure 2, bottom row), being composed of densely packed collagen 1 positive fibers. ICL1 and the IOL segments also share a taut fiber arrangement compared to the SUS and the posterior ICLs. ICL1 also has defined ovoid shape, especially in its attachment regions, which is similar to the Weberian ligaments rather than the other intercostals, which are more flattened in appearance.

**Figure 2:**
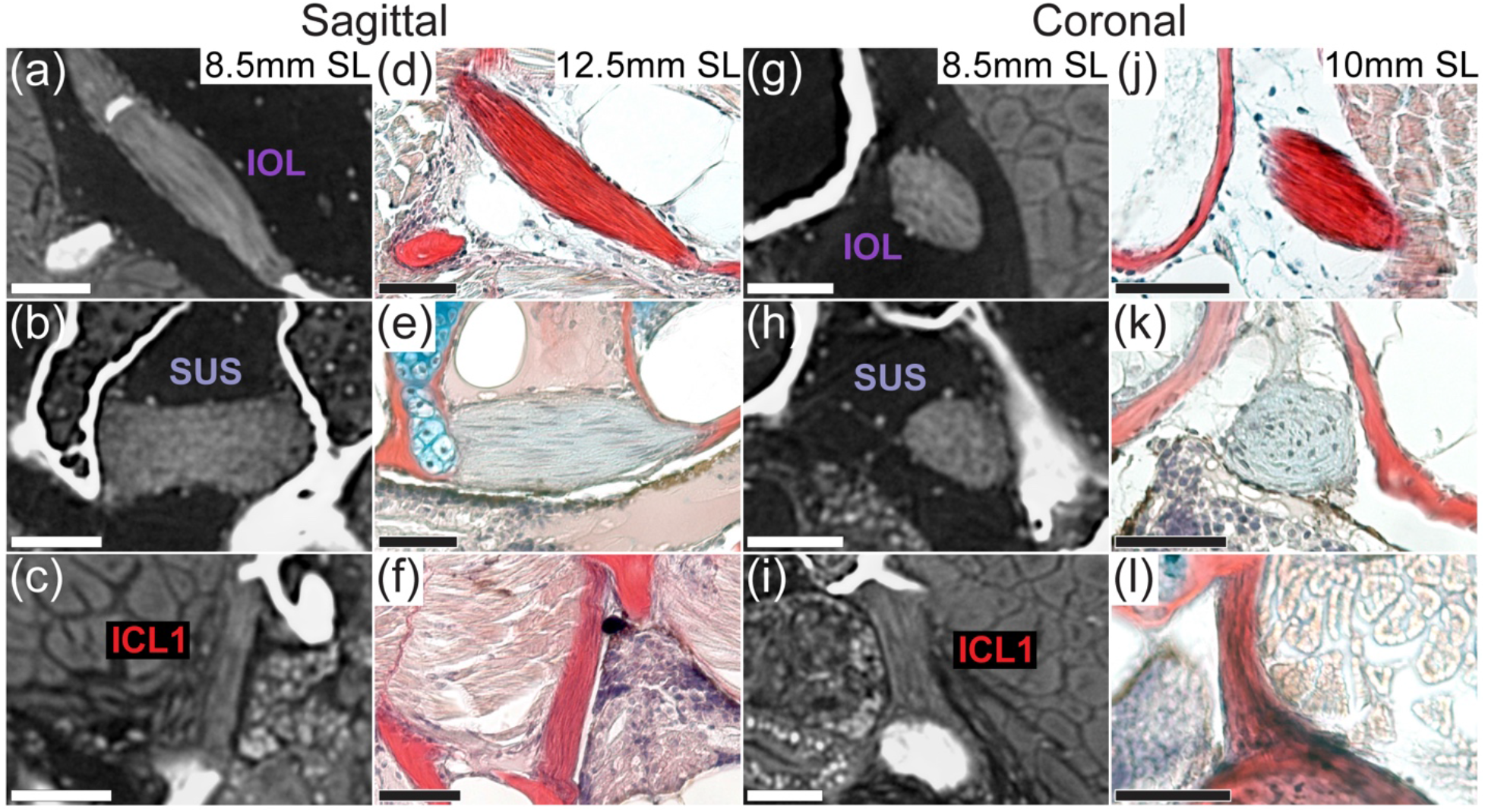
Comparison of sagittal (a-f) and coronal (g-l) virtual and histological thin sections through the Weberian ligaments and ICL1 of zebrafish. ICL: intercostal ligament, IOL: interossicular ligament, SUS: suspensor ligament. Scale bars = 40µm.

ICL1 shows distinct differences with the more posterior ICLs, indicating a potential for slightly different function. The posterior ICLs (see Figure 3 for ICL2 and ICL3) appear to have more slack compared to ICL1. All ICLs have an origin near the proximal end of the rib head of the more posterior rib, and attach more distally (proximal quarter) to the next anteriormost rib, typically along a flange or ridge in the adult rib.

**Figure 3:**
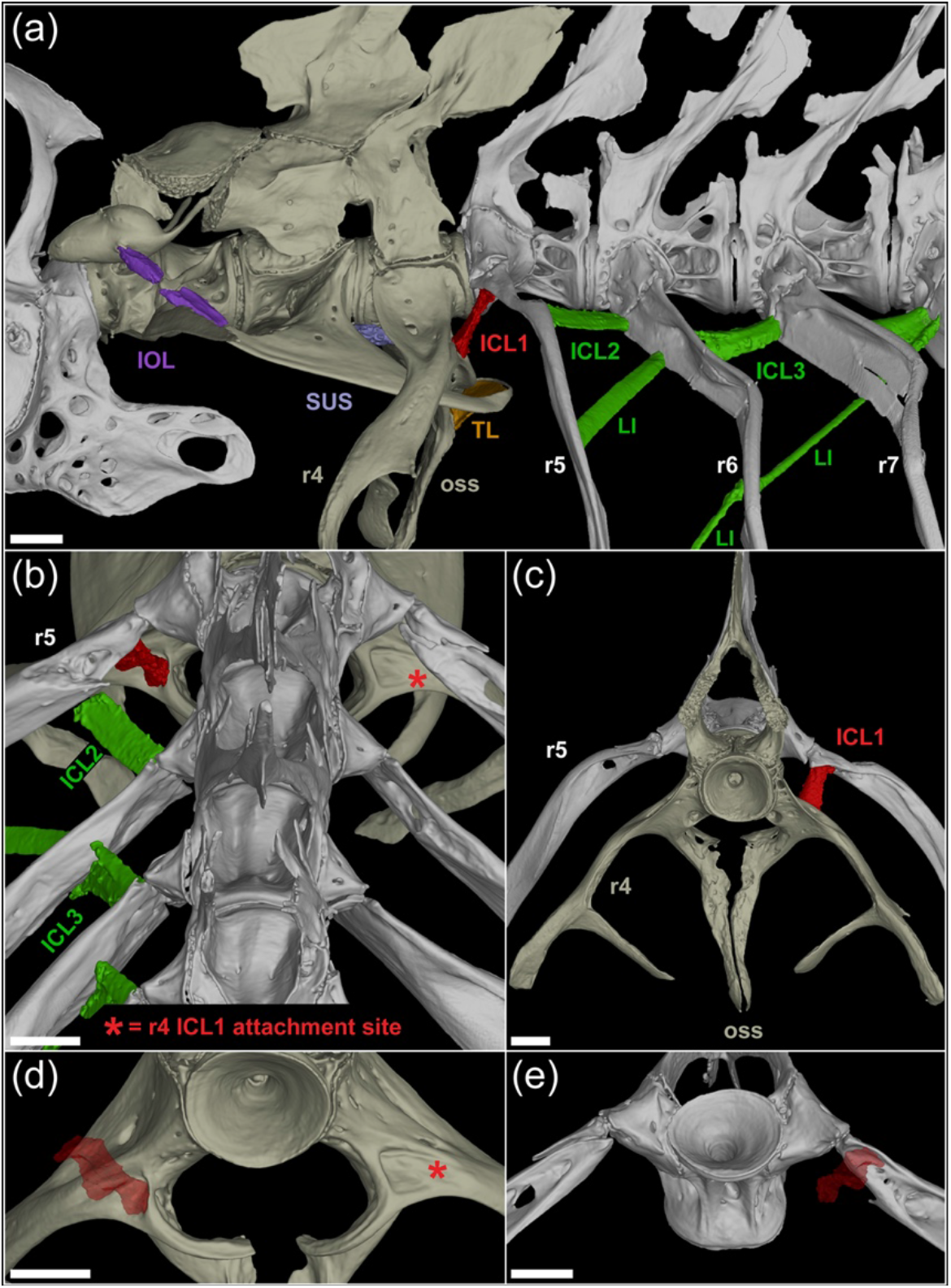
3D renderings of the intercostal ligaments of a 24mm SL zebrafish. (a) lateral view of vertebrae 1-7 including the Weberian and intercostal ligaments. (b) posterodorsal view of vertebrae 5-7 with only the left intercostal ligaments rendered. (c) anterior view of vertebrae 4 and 5 with the left ICL1 rendered. (d) posterodorsal view of the ICL1 attachment site (red asterisk) on rib/parapophysis 4. (e) anteroventral view of the ICL1 attachment site on rib 5. ICL: intercostal ligament, IOL: interossicular ligament, r: rib, SUS: suspensor ligament, oss: os suspensorium. Scale bars = 200µm.

ICL1 differs in that it connects medially, rather than laterally, and at a steep vertical angle (Figure 3) in a posterior depression marking the splitting point of rib 4 and the os suspensorium. The posterior ICLs are dorsal-ventrally flattened and oriented primarily in the anterior-posterior plane (projecting antero-laterally, Figure 3b), while ICL1 is more rounded, varies in width along its length (resembling an hour-glass shape), and is oriented in the dorsal-ventral plane (projecting ventro-medially, Figure 3b-e). This shift in direction aligns it with the os suspensorium, suggesting potential activation upon os suspensorium motion.

### 3.2 Ontogenetic development of Weberian ligaments

The first indication of ligamentous development is at 5.5 mm SL, with mesenchymal condensations (fibroblasts) near the condensing cartilage elements in the relative positions of the interossicular ligaments and the suspensor ligament, but not the triple ligament or intercostals (Figure 4a-b). At 6.0 mm SL, definitive staining of collagen fibers is present for the interossicular ligaments as well as diffuse connective tissue staining for the suspensor ligament (Data not shown; Bird et al. 2020b). At this stage, fibroblasts are seen in a large condensation in the location of the first intercostal ligament (Figure 4c). No clear development was seen for more posterior intercostal ligaments, however loose aggregations of cells can be seen. No development of the triple ligament was noted.

**Figure 4:**
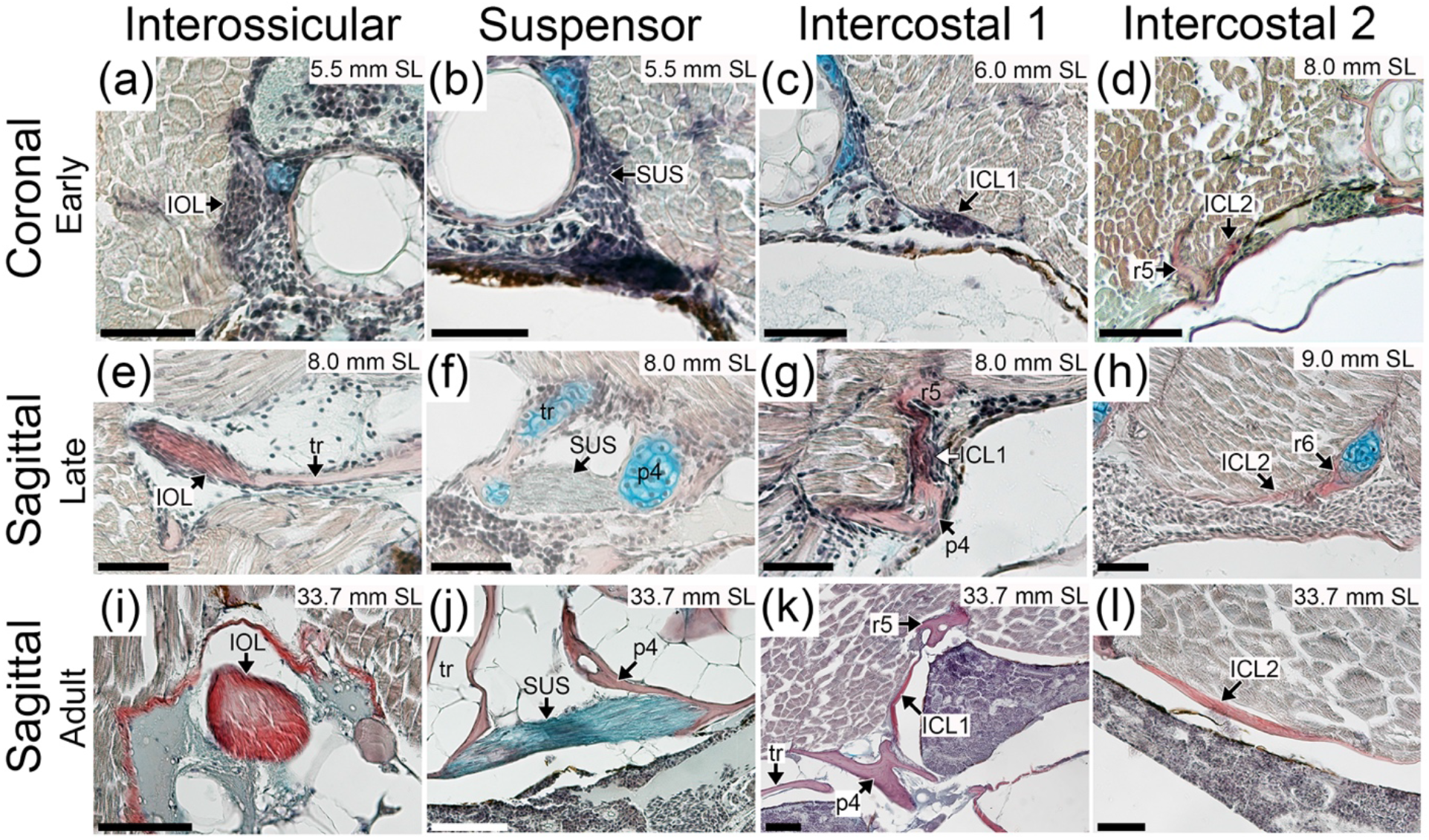
Comparative development of Weberian and intercostal ligaments in zebrafish. (a-d) in coronal plane, (e-l) in sagittal plane (anterior to the left and dorsal to the top in all sagittal images). ICL: intercostal ligament, IOL: interossicular ligament, p4: parapophysis 4, r: rib, SUS: suspensor ligament, tr: tripus. Scale bars = 40µm.

Development of the ligaments continues rapidly, with fiber deposition in ICL1 beginning at 6.5 mm SL, as well as continued growth of the interossicular and suspensor ligaments, which are composed of more fibroblasts than fibers. A small condensation is found in the presumptive location of the triple ligament (Figure 5a). By 7.0 mm SL, ICL1 is increasing in size and fiber content, while clear initial fiber staining can be seen for the triple ligament (Figure 5b) and posterior intercostal ligaments, as well initial condensations for the more posterior intercostal ligaments. The IOLs and suspensor ligament continue to increase in fiber content, with now reaching approximately a 50-50 split in cells versus fibers in their composition (Data not shown; Bird et al. 2020b).

**Figure 5:**
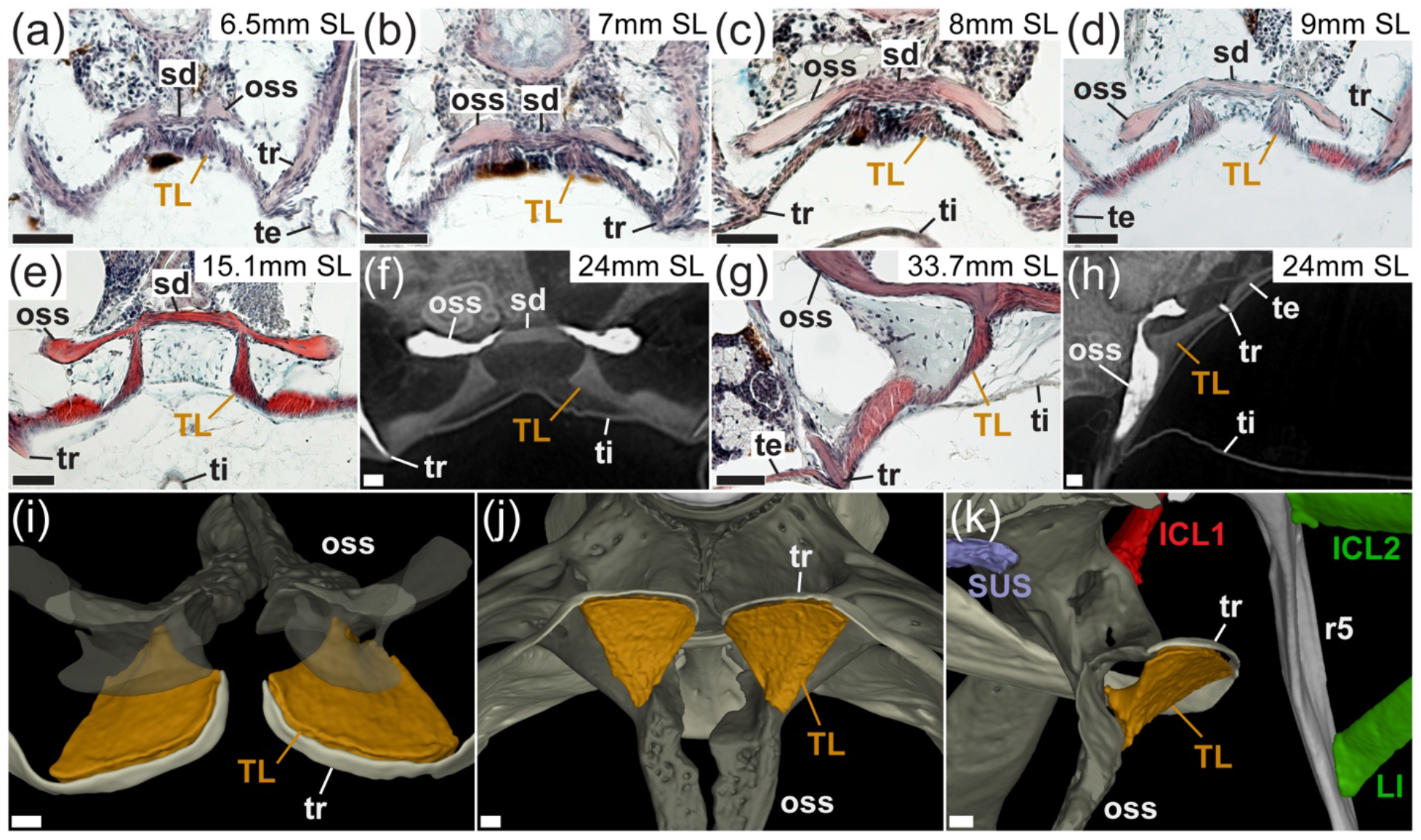
Comparative development (a-h) and reconstructed adult morphology (i-k) of the triple ligament. (a-g) in horizontal plane (anterior to the top), (h) in sagittal plane (anterior to the left and dorsal to the top), (i) anterodorsal view of the triple ligament with partially transparent os suspensorium (j) posteroventral view of the triple ligament, (k) medial view of the triple ligament. ICL: intercostal ligament (number denotes order starting from rib 4 to rib 5), LI: long intercostal, oss: os suspensorium, r: rib (number denotes vertebral level), te: tunica externa, ti: tunica interna, TL: triple ligament, tr: transformator process of the tripus, sd: sydesmosis SUS: suspensor ligament. Scale bars = 40µm.

By 8 mm SL, both ICL1 and the suspensor ligament continue growing (Figure 4f-g), ICL1 now with substantial cellularity. The IOL ligaments have transitioned now to a majority of fibers rather than fibroblasts, now densely packed with collagen-positive fibers (Figure 4e). Both the posterior ICLs (Figure 4d) and the triple ligament (Figure 5c) have become more distinct, but significantly smaller and less fiber dense compared to the more anterior ligaments. By 9 mm SL, all ligaments continue getting larger, with ICL1 and the IOLs increasing in fiber clarity. Definitive posterior ICLs are now clearly identified and gaining fiber density (Figure 4h). The suspensor ligament maintains a high level of cellularity, and its fibers are not as distinct, due to the difference in composition. Both the triple ligament (Figure 5d) and posterior intercostals have clearly defined fibers (less robust than anterior ligaments), and have taken more of an adult shape.

By 10 mm SL, all ligaments have reached near adult morphology, with shape, composition, fiber density, and cellularity approximating the adult form. Between 10-15 mm SL, no significant changes were seen in the ligaments, rather growth and size change relative to overall body growth, which continued through adult stages (Figure 4i-l, 5e-k)).

## 4. DISCUSSION

### 4.1 Intercostal Ligament 1 as a Weberian ligament, and Vertebra 5 as a transitional vertebra

ICL1 shares similarities with the IOL, and marked differences from other ICLs that suggest it should be included within the group of Weberian ligaments. The most critical evidence lies in its attachment to vertebra 4. The taut, tubular ICL1 originates from a socket in rib 5, and inserts in a groove/depression in parapophysis 4, an arrangement not seen in any other ICL. The switch from anterior-posterior orientation to dorsal-ventral likely allows for parapophysis 4 (and its distal projections) to be strongly anchored to rib 5, potentially minimizing non-optimal motion of the structure that would otherwise reduce Weberian apparatus sensitivity due to unfocused os suspensorium motion.

ICL1 is not alone in its differences with posterior elements, vertebra 5 as a whole also varies significantly from posterior rib-bearing vertebrae. Parapophysis 5 is substantially more dorsally positioned compared to those of more posterior vertebrae.

Neural arch 5 is often taller (more dorsal height) and narrower than posterior neural arches. Importantly, rib 5 varies substantially compared to other ribs, having a shorter and more rounded head (to match the parapophysis articulation site), and is more slender along its entire length (other ribs tend to have broad, flat plates in their proximal quarter).

In a comparative perscpective, the concept of vertebra 5 as a Weberian vertebra is not novel or surprising. Particularly in many catfish species (Siluriformes), the incorporation of vertebra 5 and its structures into the Weberian apparatus structure is well established. In the developing structure of *Clarius gariepinus*, the basiventrals (parapophyses) of vertebra 5 are incorporated into the broader Weberian structure (Radermaker et al., 1989). Incorporation of the fifth parapophysis into a thin sheet (plate) overlying the swim bladder to the skin, or expanded to a near capsule, was noted in primitive Siluriforms (including *Clarias*) by Alexander (1964b). These species, as well as *Plecostomus*, often incorporate centrum 5 into their fused *complex centrum* (Alexander, 1964b). Chardon et al. (2003) also described various contributions of elements of the fifth vertebra to the overall Weberian structure in catfishes. While zebrafish (and other Cypriniformes) do not incorporate vertebra 5 into the Weberian apparatus to the extent of siluriform species, taken broadly, it is clear that vertebra 5 should, at a minimum, be considered a transitional vertebra between classically defined Weberian vertebra and posterior rib-bearing vertebrae, and the modifications suggest this vertebral level is responsive to the Weberian developmental module.

### 4.2 Possible functional roles of the V5 intercostal ligament and updated model

There are several anatomical features that both distinguish ICL1 from more posterior intercostal ligaments, and point strongly to a unique role related to the function of the Weberian apparatus. First, the shape differences between ICL1 and more posterior ICLs are distinct, with ICL1 being more cylindrical, while posterior ICLs are more broad and flat (Figure 3). Second, ICL1 has much different orientation, being nearly vertical in the dorsal-ventral axis, while posterior ICLs are predominantly flattened along the horizontal axis. Third, while posterior ICLs attach between ribs broadly onto small flanges in the ribs, ICL1 attaches in defined, deep depressions in parapophysis 4 and rib 5 (Figure 3b-e), likely allowing for more directed and focused action. These differences clearly point to a shift in function for ICL1. When paired with the slight medial direction at its parapophysis 4 attachment relative to its origin, this angle puts ICL1 in alignment with the os suspensorium, making it likely to respond to movement of the os suspensorium resulting from swim bladder pulsation. Given that parapophysis 4 forms a synchondrosis with centrum 4 (Bird and Mabee 2003), there is some theoretical rotation possible at that articulation. It is plausible that ICL1 is acting to limit unnecessary rotation of parapophysis 4, potentially in multiple planes, to prevent vibrational signal loss due to spurious movement of the surrounding skeletal structures of the Weberian apparatus, thereby helping to direct movement along the ossicle chain and increased sensitivity with minimal signal loss.

### 4.4 Future directions

This and other recent studies (Bird et al. 2020a, b) underscore the need for increased focus on the ligamentous components of the Weberian apparatus. For a truly comprehensive understanding of the Weberian apparatus and how it functions, more integrative and holistic approaches are needed that include all regions and all ligaments. We outline several areas of needed future research below with focus on the ligaments.

#### Molecular/Functional

In order to create realistic and testable hypotheses regarding how each of the Weberian ligaments, including ICL1, function independently (as well as collectively), a comprehensive molecular marker characterization and fiber composition studies are needed. Known fibers in the various ligaments include collagens, elastin, and others. Immunohistochemical and in situ hybridization analyses in these ligaments would confirm the molecular markers for different fiber types. In addition, ratios of the fiber content in the various ligaments would allow for hypotheses regarding tensile strength, which can then be related to overall function.

#### Comparative

Given the lack of ICL1 in the description of the Weberian apparatus in cypriniform fishes, a reexamination vertebrae 4-5 is needed to properly assess both the presence of this ligament across species, as well as the structural connectivity of ICL1 with rib 5 and parapophysis 4. Primary focus should start with species possessing the ‘Open’ Weberian morphology (Bird et al. 2020a), which are most likely to have a structural similarity to the zebrafish. Surveys should then extend to those with anterior plates, such as gyrinocheilids, catostomids, and some botids. A much broader survey of the fate of ICL1 could then be undertaken on encapsulated species (cobitids, nemacheilids, balatorids, etc.), as well as other otophysan clades such as Characiformes to determine the overall incorporation of ICL1 into the Weberian apparatus in disparate species.

#### Rib 4 and the Os Suspensorium

The precise role of the os suspensorium in the function of the Weberian apparatus has remained elusive. In the zebrafish, the os suspensorium is deeply embedded within the tissue surrounding the swim bladder and is connected to the tripus and swim bladder tunica via the triple ligament (Bird et al. 2020b, Figure 5). In addition, parapophysis 4, which bears the os suspensorium, is connected to the articular process of the tripus via the suspensor ligament. Still less is known about a possible role of rib 4. Given that parapophysis 4 articulates with centrum 4 via a synchondrosis, potential mobility is present at this joint. One possible role of rib 4 would be to provide balance to the entire vertebra 4 structure, and the added mass could provide inertial resistance to excessive motion of the os suspensorium, however this and other hypotheses regarding the functions of the os suspensorium and rib 4 require empirical testing.

## ACKNOWLEDGEMENTS

The study was supported by the Swedish Research Council, Grant number: 2022-04988 and Magnus Bergvalls Stiftelse awarded to TH; and by University of Northern Iowa Department of Biology Research Funds awarded to NCB. The beamtime was received as part of proposal LS3021 accepted at the European Synchrotron Radiation Facility – Extremely Brilliant Source (ESRF-EBS) in France on the beamline BM05. We acknowledge Dr. Paul Tafforeau and Dr. Kathleen N. Dollman for synchrotron data aquisition and reconstruction. We thank Assoc. Prof. Sophie Sanchez at Uppsala University for providing access to VGStudio MAX workstation. Open access publishing facilitated by Uppsala University, as part of the Wiley - Bibsam agreement.

## 5. AUTHOR CONTRIBUTIONS

JL and TH contributed to the conceptualisation of the study. All authors participated in data collection, data analysis, interpretation, and drafting of the original manuscript.

